# HORDCOIN: A Software Library for Higher Order Connected Information and Entropic Constraints Approximation

**DOI:** 10.64898/2026.02.08.704639

**Authors:** Giulio Tani Raffaelli, Jakub Kislinger, Tomáš Kroupa, Jaroslav Hlinka

**Affiliations:** Department of Data-Analysis, Ghent University, 1 Henri Dunantlaan, Ghent, B-9000, Belgium; Institute of Computer Science, Czech Academy of Sciences, Pod Vodárenskou věží 2, Prague, 182 00, Czech Republic; Faculty of Electrical Engineering, Czech Technical University in Prague, Technická 2, Prague, 166 27, Czech Republic; National Institute of Mental Health, Topolová 748, Klecany, 250 67, Czech Republic

**Author notes:** G. Tani Raffaelli and J. Kislinger contributed equally to this work.

**Keywords:** connected information, maximum entropy, higher-order interactions, information theory, neuroimaging, neurophysiology

## Abstract

**Background and objective:** Quantifying higher-order statistical dependencies in multivariate biomedical data is essential for understanding collective dynamics in complex systems such as neuronal populations. The connected information framework provides a principled decomposition of the total information content into contributions from interactions of increasing order. However, its application has been limited by the computational complexity of conventional maximum entropy formulations. In this work, we present a generalised formulation of connected information based on maximum entropy problems constrained by entropic quantities.

**Methods:** The entropic-constraint approach, contrasting with the original constraints based on marginals or moments, transforms the original nonconvex optimisation into a tractable linear program defined over polymatroid cones. This simplification enables efficient, robust estimation even under undersampling conditions.

**Results:** We present theoretical foundations, algorithmic implementation, and validation through numerical experiments and real-world data. Applications to symbolic sequences, large-scale neuronal recordings, and DNA sequences demonstrate that the proposed method accurately detects higher-order interactions and remains stable even with limited data.

**Conclusions:** The accompanying open-source software library, HORDCOIN (Higher ORDer COnnected INformation), provides user-friendly tools for computing connected information using both marginal- and entropy-based formulations. Overall, this work bridges the gap between abstract information-theoretic measures and practical biomedical data analysis, enabling scalable investigation of higher-order dependencies in neurophysiological and other complex biological systems such as the genome.

## 1. Introduction

Understanding and characterizing complex systems from empirical multivariate observations or recorded time series of activity represents a key challenge across many scientific disciplines. Such systems often comprise numerous interacting components whose collective dynamics give rise to emergent behavior that cannot be predicted from individual elements alone. A quantitative description of these interactions is essential for uncovering the underlying mechanisms and functional organization.

This challenge is particularly prominent in neuroscience, where the brain’s activity reflects interactions within large neuronal populations recorded through electrophysiological or neuroimaging techniques [10, 27, 26]. Similar issues arise analyzing other biomedical signals, as well as in other domains such as climate science, economics, or ecology, where multiple interdependent subsystems jointly shape complex global dynamics [6, 7]. In all these fields, robust methods to infer and quantify interaction structure from multivariate data are crucial for advancing both understanding and predictive modeling.

A widely adopted strategy to describe such dependencies is the network approach, where system components are represented as nodes and their statistical relationships as links [10, 27]. This formalism enables both visualization and quantitative analysis of the interaction architecture and has become a cornerstone of research in areas such as brain connectivity, gene regulatory networks, and climate interaction mapping.

However, most existing approaches rely on pairwise measures, such as correlation or mutual information. While these information measures capture direct interactions between pairs of variables, they inherently neglect higherorder dependencies involving three or more components. As a result, network models constrained to pairwise interactions may fail to account for collective or synergistic effects that emerge only at the multivariate level [28, 37, 6].

To overcome these limitations, methods capable of identifying and quantifying higher-order interactions have been proposed. These approaches aim to reveal dependencies that cannot be reduced to lower-order effects and can thus provide deeper insight into system organization, including the identification of synergistic and redundant information sharing among multiple components [7, 26].

A fundamental first step is to estimate the overall contribution of higher-order interactions to the structure of the system’s activity. This requires decomposing the total information content into components corresponding to interactions of increasing order [29, 1, 20].

A principled framework for such decomposition was introduced by Schneidman et al. [29], based on the information-geometric perspective of Amari [1]. The key concept of *connected information* provides a systematic way to quantify the informational contribution of interactions of different orders within a multivariate distribution.

In this framework, the entropy of the observed joint distribution is compared with the maximum entropy achievable by a model constrained to reproduce marginal distributions up to order *k*. The difference between the two reflects the amount of information carried by interactions beyond order *k*.

The difference between the maximum entropies when fixing marginals up to orders *k* and *k* + 1 defines the connected information of order *k*, quantifying how much additional dependency structure emerges when moving from *k*-wise to (*k* + 1)-wise interactions.

Computing these quantities is, however, computationally demanding, since finding the maximum-entropy distribution under fixed marginal constraints requires highdimensional optimization. Schneidman et al. therefore proposed a tractable approximation based on constraining the statistical moments rather than the full marginals [29]. While moment-based formulations simplify computation, they remain theoretically limited and sensitive to sampling. To address this, Martin et al. [22, 23] introduced an alternative formulation in which the maximum entropy is computed under constraints on marginal *entropies* up to order *k*. This entropy-based approach allows the problem to be expressed as a linear optimization task over information-diagram atoms, leading to a fast and distribution-free estimate of connected information.

In practice, the entropy-constrained connected information can be efficiently approximated using linear programming, providing a computationally scalable and robust estimator even in undersampled regimes often encountered in neural and other biological data [23, 25].

Although these measures have been successfully used to study the collective dynamics of neuronal populations and resting-state brain networks [28, 22, 23], their technical and computational complexity has so far limited wider adoption in the biomedical community.

Here, we address this bottleneck by introducing HORDCOIN, (Higher-ORDer COnnected INformation, Hordcoin.jl), a Julia package that implements several variants of connected information estimation, including both marginal- and entropy-based formulations [32]. HORDCOIN provides an efficient, user-friendly framework for quantifying higher-order dependencies in multivariate data, facilitating its application to neurophysiological, neuroimaging, and other complex biomedical datasets such as the genome. For additional ease of use, we released an official Python interface pyHordcoin allowing to access the functionalities and speed of Hordcoin.jl without requiring familiarity with Julia [33]. This paper presents the theoretical background, computational implementation, and illustrative applications of the HORDCOIN library.

In Section 2, we summarize the mathematical preliminaries underlying the connected information framework. Section 3 defines the connected information formally and provides illustrative examples. Section 4 describes the entropy-constrained formulation and its approximation via linear programming.

Section 5 reports numerical experiments validating the performance and efficiency of the implemented algorithms. Section 6 demonstrates their use in practical settings, including higher-order interaction analysis in symbolic data (English and German texts), in large-scale neuronal spike recordings from public datasets, and DNA sequences. A brief discussion and conclusion follow in Section 7.

### 2. Preliminaries

In this section, we introduce key concepts and results from information theory. We adopt notation analogous to that originally developed for analyzing interaction spaces of random variables; see [13] or [3, Chapter 2.9.1]. Let *N* = {1, …, *n*} be the index set for some integer *n ≥* 1. For each *i* ∈ *N* we consider a nonempty finite sample space 𝒳_*i*_ ⊂ ℝ. For any nonempty *A* ⊆ *N* let 𝒳_*A*_ = ×_*i*_._∈*A*_ 𝒳_*i*_.For simplicity, we write 𝒳= 𝒳_*N*_ whenever the index set *N* is clear from context. Throughout the paper we consider a discrete random vector **X** = (*X*_*i*_)_*i* ∈*N*_ with the finite sample space 𝒳 and a joint probability distribution. Specifically, function *p*: 𝒳 → [0, 1] satisfies ∑ _x∈𝒳_*p*(**x**) = 1. Let Δ be the set of all probability distributions on 𝒳, which coincides with the (|𝒳 | − 1)-dimensional standard simplex. For any nonempty *A* ⊆ *N*, let **X**_*A*_:= (*X*_*i*_)_*i*∈*A*_ and Δ_*A*_ be the set of all probability distributions on *X*_*A*_. Observe that **X**_*N*_ = **X** and Δ_*N*_ = Δ. For any probability distribution *p* ∈ Δ of **X**, the marginal probability distribution *p*_*A*_ ∈ Δ_*A*_ of random vector **X**_*A*_ is given by

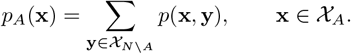

The probability distribution *p*_*A*_ is also called the *A-marginal of p*. In case *A* = {*i*}, we also write *p*_*i*_ = *p* _{*i*}_ and call *p*_*i*_ the *i*-marginal of *p*. When variables *X*_1_, …, *X*_*n*_ are mutually independent, their joint distribution is the *product distribution p*^∗^ ∈ Δ defined by

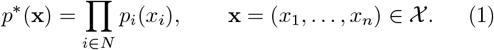

Throughout the paper, we make use of several information-theoretic functionals [3, 12]. For any *p*_*A*_ ∈ Δ_*A*_, the set spt *p*_*A*_ = {**x** ∈ 𝒳_*A*_ *p*_*A*_(**x**) *>* 0} is the *support* of *p*_*A*_. The *(Shannon) entropy* of **X**_*A*_ is

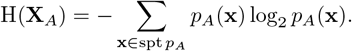

Since the functional H depends only on the values of *p*_*A*_, we will often write H(*p*_*A*_) in place of H(**X**_*A*_). The *KullbackLeibler divergence of p* ∈ Δ *from q* ∈ Δ such that spt *p*⊆ spt *q* is

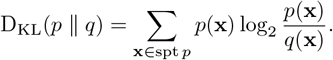

The *multi-information* [31] (or, equivalently, *total correlation* [36]) of **X** defined by

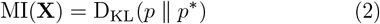

is a general measure of non-independence among variables in **X**. The special case of two random variables, *n* = 2, yields the *mutual information* between *X*_1_ and *X*_2_,

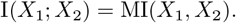

The basic identity

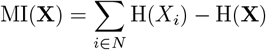

can be derived directly from the definition. The mutual independence of random variables *X*_1_, …, *X*_*n*_ is characterized by MI(**X**) = 0. Conversely, MI(**X**) *>* 0 indicates dependence among the variables. However, it remains unclear whether this dependence arises from interactions involving all variables *X*_1_, …, *X*_*n*_ collectively or only specific subsets of them. For example, let *p* be a joint distribution of (*X*_1_, *X*_2_, *X*_3_) factorizing as *p* = *p* _{1}_ *p* _{2,3}_. Then MI(*X*_1_, *X*_2_, *X*_3_) *>* 0 although there is no three-way interaction.

This observation motivates the search for decomposition of MI(**X**) into nonnegative components [29, 2, 4], where each summand captures interactions of a particular order (e.g., pairwise, triple-wise) separately. In Section 3, we concentrate on a specific decomposition grounded in the concept of connected information.

### 3. Connected Information

This section presents the construction of connected information under entropic constraints (Definition 1), as originally introduced in [23]. We begin by formulating and analyzing the maximum entropy problem with entropic constraints (3), since solving this problem directly yields the values of the connected information.

Let *p* ∈ Δ be a probability distribution of a random vector **X** = (*X*_1_, …, *X*_*n*_) on the finite sample space 𝒳. For each *k* ∈ *N* = {1, …, *n*}, let

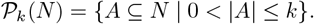

If the set *N* is clear from context, we write simply 𝒫_*k*_ = 𝒫_*k*_(*N*). We will deal with the following family of probability distributions:

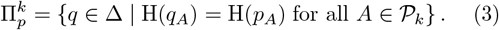

The set 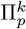 contains all joint distributions *q* of **X** that preserve the entropic structure of every subset of variables of size at most *k*. This approach, introduced in [22], proposes an alternative formulation of the maximum entropy problem [29, 2], where all *k*th-order marginals of the joint distribution *q* ∈ Δ are constrained to match the corresponding marginals of the input distribution *p*, specifically:

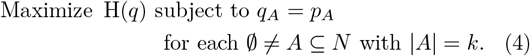

For a detailed survey of maximum entropy problems with given marginals or moments, see [16].

As a side note, note that, in general, imposing only the 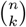 constraints of the form H(*q*_*A*_) = H(*p*_*A*_) for all *A* ⊆ *N* with |*A*| = *k* does not suffice to guarantee that 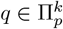. Equivalently, the entropy constraints on all lower-dimensional marginals *q*_*A*_ with 0 *<* |*A*| *< k* cannot be omitted without altering the meaning of the definition (3); see the second part of Example 2. This stands in sharp contrast to the marginal constraints in problem (4), since the identities *q*_*A*_ = *p*_*A*_ for ∅ ≠ *A* ⊆ *N* imply *q*_*B*_ = *p*_*B*_ for all ∅≠*B* ⊆ *A*.

These important inclusions follow directly from (3):

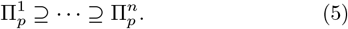

For each *k* ∈ *N*, the set 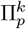 is nonempty (since 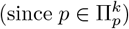) and compact, as the entropy functional is continuous. However, as illustrated by the following examples, 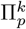 is generally not a convex set.

#### Example 1

(*n* = *k* = 1). *Let X be a binary random variable over* 𝒳 = {0, 1} *with probability distribution p such that* 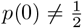. *Then* 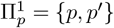, *where p*^*′*^ *is a probability distribution defined by p*^*′*^(0) = *p*(1) *and p*^*′*^(1) = *p*(0).

#### Example 2

(*n* = 2 and *k* = 1, 2). *Let 𝒳*_1_ = 𝒳_2_ = 0, 1. *We consider the probability distribution p of binary random vector* (*X*_1_, *X*_2_) *specified in Table 1*.

**Table 1:**
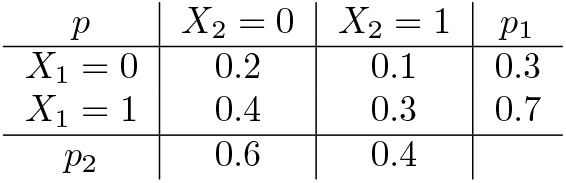
Probability distribution *p* and its marginals *p*_1_ and *p*_2_.

*The entropies of marginals are* H(*p*_1_) ≐ 0.8813 *and* H(*p*_2_) *≐* 0.9710. *Since X*_1_ *and X*_2_ *are binary variables, the constraints* H(*q*_1_) = H(*p*_1_) *and* H(*q*_2_) = H(*p*_2_) *imply that the one-dimensional marginal of any* 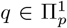 *differs from the corresponding one-dimensional marginal of p only by the flip. This implies that there are four possible cases, which are displayed in Tables 2a–2d, each representing a distinct class of probability distributions with specified one-dimensional marginals. As a result, the set* 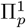 *is the union of four pairwise disjoint line segments within the 3-dimensional standard simplex* Δ.

**Table 2:**
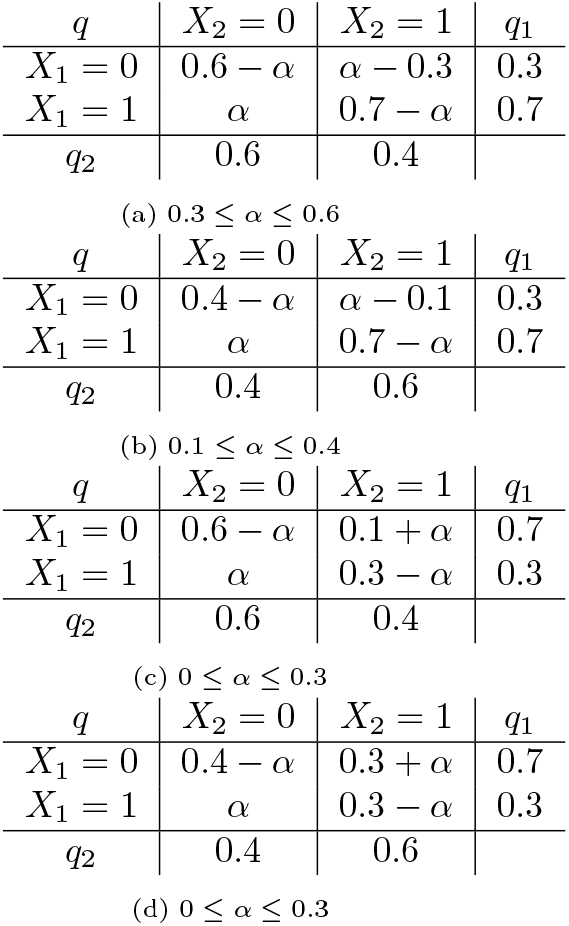
Probability distributions *q* preserving the marginal distributions.

*By the definition*,

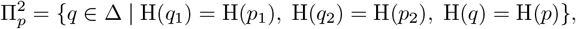

*which means that* 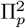 *is the intersection of* 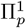*with the level set of entropy given by*

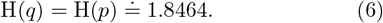

*Specifically*, 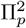 *is a finite set comprising 8 probability distributions, with each of the 4 pairs of distributions corresponding to one of Tables 2a–2d based on the constraint* (6); *see Tables 3a–3d. In each of these tables, the probability distributions* 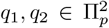 *are associated with α*_1_ *and α*_2_, *respectively*.

**Table 3:**
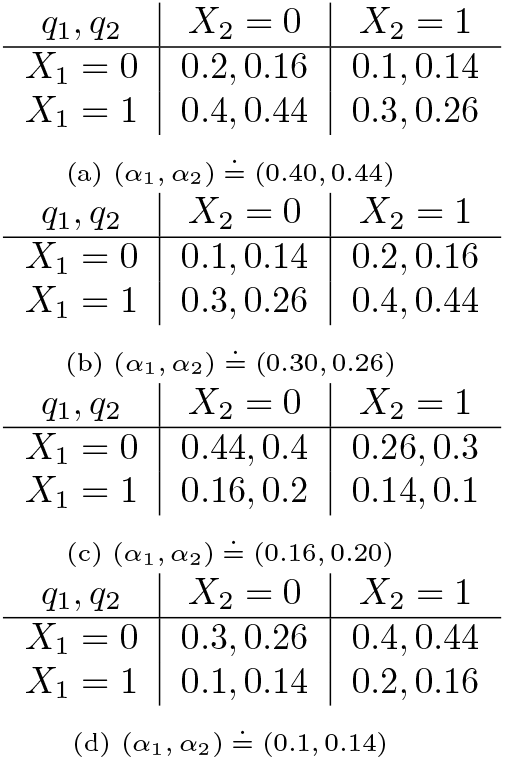
Probability distributions *q* preserving the marginal distributions.

Computing the connected information (see the Definition 1 below) involves solving the following nonconvex optimization problem for every *p* ∈ Δ and each *k* = 1, …, *n*:

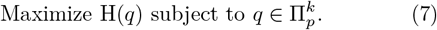

Optimization problem (7) is a relaxation of the classic maximum entropy problem (4). Indeed, the feasible set of (4) is included in the feasible set of (7). This also means that the optimal value of (7) is an upper bound on (4). Although problem (7) introduces 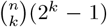 entropic constraints and is nonconvex—offering no immediate computational advantage over the convex formulation (4) with only 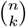 marginal constraints—we will show in Section 4 that (7) admits computationally tractable (linear) approximations.

Since the entropy functional H is continuous and the feasible set 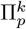 is nonempty and compact, the problem (7) is guaranteed to have at least one global maximizer. Let 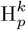 denote the optimal value of (7),

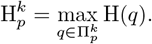

In general, the globally optimal solution to (7) need not be unique and computing the global maxima is highly nontrivial.

We will derive analytical solutions for several fundamental instances of (7), illustrating the structure of the problem and providing insight into its key properties. If *k* = *n*, then

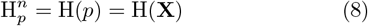

because H(*q*) = H(*p*) is one of the constraints, fixing the objective function in (7) to the constant H(*p*). For *k* = 1 and arbitrary *n*, the solution to (7) is the product distribution. Every *q* ∈ Δ satisfies the inequality H(*q*) ≤ ∑ _*i*∈*N*_ H(*q*_*i*_), where *q*_1_, …, *q*_*n*_ are the one-dimensional marginals of *q*. The equality H(*q*) = ∑ _*i*∈*N*_ H(*q*_*i*_) holds if and only if *q* = *q*^∗^, where *q*^∗^ denotes the product distribution (1). By definition, 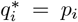 for each *i* ∈ *N*, which implies 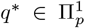. Therefore, *q*^∗^ is the maximizer of problem (7) when *k* = 1, and the optimal value is given by

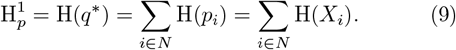

The analytic computation of 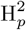 for three random variables was given in [23]. For completeness, we re-derive this result below using an elementary argument.

#### Example 3

(*n* = 3 and *k* = 2). *In this setting the problem* (7) *is to maximize* H(*q*) *under the entropic constraints* H(*q*_*i*_) = H(*X*_*i*_) *for each i* = 1, 2, 3 *and* H(*q*_{*i,j*}_) = H(*X*_*i*_, *X*_*j*_) *for all i, j* ∈ {1, 2, 3} *with i < j, where* 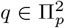 *is a probability distribution of* **X** = (*X*_1_, *X*_2_, *X*_3_).

*By submodularity (See Section 4*.*1 for details) of entropy we obtain the upper bound on the objective:*

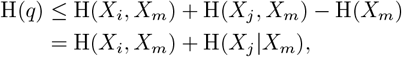

*where the indices i, j, m* ∈ {1, 2, 3} *are selected so that*

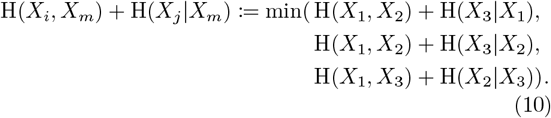

*We will find* 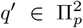 *such that* H(*q*^*′*^) = H(*X*_*i*_, *X*_*m*_) + H(*X*_*j*_ |*X*_*m*_).

*The idea is to construct q*^*′*^ *so that X*_*i*_ *and X*_*j*_ *become conditionally independent given X*_*m*_. *Specifically, define*

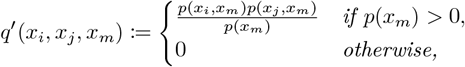

*for all x*_*i*_ ∈ 𝒳 _*i*_, *x*_*j*_ ∈ 𝒳_*j*_, *x*_*m*_ ∈ 𝒳_*m*_. *By the definition, q*^*′*^ *has the same marginals of order* 2 *as p, and hence* 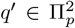. *Moreover, X*_*i*_ *and X*_*j*_ *are conditionally independent given X*_*m*_ *under q*^*′*^, *so the conditional mutual information of X*_*i*_ *and X*_*j*_ *given X*_*m*_ *with respect to q*^*′*^ *vanishes*,

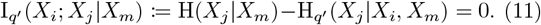

*Then*

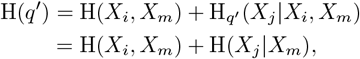

*where the first equality follows from the chain rule for entropy and the second equality from* (11). *In conclusion*,

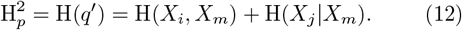

The notion of connected information in the context of entropic constraints was first introduced in [23].

#### Definition 1.

Let *p* ∈ Δ be a probability distribution of random vector **X** = (*X*_1_, …, *X*_*n*_). The *connected information* of order *k* = 2, …, *n* is

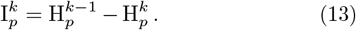

It follows from the inclusions (5) and identities (8), (9) that

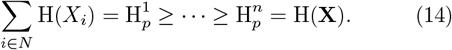

This implies that the connected information is always nonnegative, 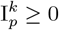. Aggregating the connected information across all interaction orders results in

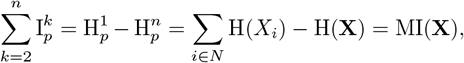

where MI(**X**) is the multi-information (2). The number 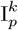 indicates the level of *k*-order stochastic interactions among random variables *X*_1_, …, *X*_*n*_ with the distribution *p*. In the trivial case of independence, no interactions exist, as this implies *p* = *p*^∗^, resulting in 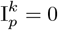 for each *k* = 2, …, *n*. The connected information admits a closed-form analytic expression only in a few simple cases.

#### Example 4

(*n* = *k* = 2). *We get*

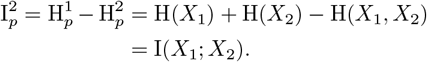

#### Example 5

(*n* = 3 and *k* = 2). *It follows from* (12) *that*

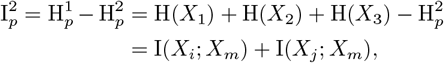

*where i, j, m* ∈ {1, 2, 3} *are such that* (10) *holds true*.

The following well-known example demonstrates that a binary three-dimensional random vector can exhibit genuine third-order interactions while having no second-order interactions.

#### Example 6.

*Let 𝒳*_1_ = 𝒳_2_ = 𝒳_3_ = 0, 1. *Assume that variables X*_1_ *and X*_2_ *are uniform, independent, and define X*_3_ = XOR(*X*_1_, *X*_2_). *The joint probability distribution p is given by Table 4*.

**Table 4:** *X*_3_ = XOR(*X*_1_, *X*_2_)

| $X_1$ | $X_2$ | $X_3$ | $p$ |
| --- | --- | --- | --- |
| 0 | 0 | 0 | $1/4$ |
| 0 | 1 | 1 | $1/4$ |
| 1 | 0 | 1 | $1/4$ |
| 1 | 1 | 0 | $1/4$ |

*Then* I(*X*_1_; *X*_2_) = I(*X*_1_; *X*_3_) = I(*X*_2_; *X*_3_) = 0, *which implies* 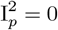 *by Example 5. Further, by* (12),

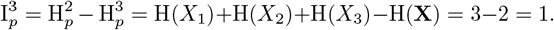

### 4. Approximation of Connected Information

The feasible set 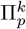 in the maximum entropy problem with entropic constraints (7) is notably complex, as it is defined by the finite intersection of entropy level sets. Consider *A* ∈ 𝒫_*k*_ with | 𝒳|_*A*_ *≥* 3. According to [8, Theorem 3], no level set defined by H(*q*_*A*_) = H(*p*_*A*_) is semialgebraic (a subset of Euclidean space is *semialgebraic* if it can be described as the solution set of finitely many polynomial equations or inequalities), except in the extreme cases where H(*p*_*A*_) = 0 or H(*p*_*A*_) = log_2_ |𝒳_*A*_|. Furthermore, the same result [8, Theorem 3] implies that under these conditions, the constraints H(*q*_*A*_) ≥ H(*p*_*A*_), which form part of a straightforward convex relaxation of the nonconvex problem (7), also define a non-semialgebraic set. This lack of semialgebraicity poses a significant obstacle to the application of numerical algorithms based on semidefinite optimization and real algebraic geometry [5, 9].

Therefore, developing computationally tractable approximations to (7) is a challenging problem. Our approach starts with the observation that both the objective function and the constraints of (7) depend solely on entropy values. To leverage this, we utilize the well-known properties of entropic vectors to reformulate the original problem (7) as an optimization problem involving only variables that represent entropies. This idea was first introduced in [23]. A potential drawback of this reformulation is that the optimal solutions to the resulting problem may not correspond to any probability distribution. Nevertheless, the transformed problem will be a linear mathematical program, providing an upper bound on the optimal value of (7). In particular, we examine the outer approximation of the entropic region via the polymatroid cone, under the assumption of a fixed sample space. For additional details, refer to [38].

### 4.1. Polymatroids and entropic cone

Entropic functionals of the form H(**X**_*A*_) are special examples of polymatroids [17] when *A* ranges in all subsets of *N*. Let 𝒫 (*N*) be the set of all subsets of *N*. A *polymatroid* is a function *h*: 𝒫 (*N*) → ℝ satisfying the following properties for all *A, B* ⊆ *N*:

1. *h*(∅) = 0,
2. *h*(*A*) ≥ h(B) whenever *A* ⊇ B, (*monotonicity*)
3. *h*(*A*) + *h*(*B*) ≥ h(A ∪ B) + *h*(*A* ∩ B). (*submodularity*)

Let Γ_*n*_ be the set of all polymatroids on 𝒫 (*N*). For every *α*_1_, *α*_2_ ≥ 0 and all *h*_1_, *h*_2_ ∈ Γ_*n*_, we obtain *α*_1_*h*_1_ +*α*_2_*h*_2_ ∈ Γ_*n*_. In other words, the set Γ_*n*_ is a convex cone in the linear space of all functions 𝒫 (*N*) → ℝ. Clearly, the convex cone Γ_*n*_ is polyhedral since it is defined by finitely-many linear inequalities of the form 1.–3. It is also pointed, that is, it doesn’t contain a nontrivial linear subspace, since every *h* ∈ Γ_*n*_ is a nonnegative function. We call Γ_*n*_ a *polymatroid cone (of order n)*.

It can be shown [11, 38] that special 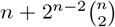 *elemental inequalities* form the irreducible representation of Γ_*n*_. Namely the monotonicity inequalities

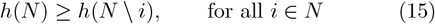

and the submodular inequalities of the form

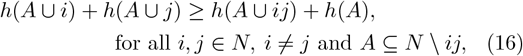

where we use the short notation *i* = *i* and *ij* = *i, j*. This means that a function *h*: 𝒫 (*N*) → ℝ is a polymatroid if, and only if, *h*(∅) = 0 and (15)–(16) hold. A *Shannon-type inequality* is any linear inequality which is a nonnegative linear combination of elemental inequalities (15)–(16).

The basic examples of polymatroids are entropic vectors. Let *p* ∈ Δ be a joint probability distribution of a discrete random vector **X** = (*X*_1_, …, *X*_*n*_). Define

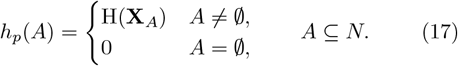

We say that *h*_*p*_: 𝒫 (*N*) → ℝ is an *entropic vector*. It follows from the basic properties of entropy (monotonicity) and conditional mutual information (submodularity) that *h*_*p*_ is a polymatroid, *h*_*p*_ ∈ Γ_*n*_. For each integer *n ≥* 1, the *entropic region (of order n)* 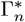 is defined as the set of all entropic vectors *h*_*p*_ induced by joint probability distributions *p* of discrete random vectors (*X*_1_, …, *X*_*n*_), where each *X*_*i*_ takes values in all possible finite sample spaces 𝒳_*i*_. Clearly, 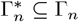 for every *n ≥* 1. We will briefly review the properties of entropic regions for *n* = 2, 3, 4; see [38, 39, 11] for further details.

- *Case n* = 2. Then 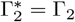.
- *Case n* = 3. The strict inclusion 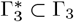 holds. However, the equality holds at least for the topological closure of 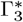, that is, 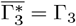.
- *Case n* = 4. It was shown that 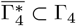 by constructing a non-Shannon type linear inequality valid for 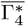 Specifically, *Zhang-Yeung inequality* for a function *h*: 𝒫 (*N*) → ℝ is a linear inequality

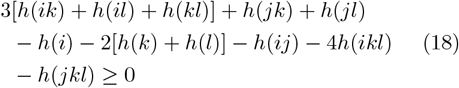

where *i, j, k, l* are different elements of *N* = {1, 2, 3, 4}. Note that there are 12 instances of this inequality each of which corresponds to the set-tuple {*k, l*}, (*i, j*). Summing up, every entropic vector in four variables satisfies all instances of linear inequality (18), and no such instance is a nonnegative linear combination of Shannon-type inequalities.

Matúš [24] established that 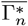 is not polyhedral for *n* ≥ 4. These findings suggest that determining whether a (2^*n*^ − 1)- dimensional real vector belongs to 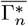 or 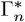 is undecidable. However, to the best of our knowledge, a formal proof of this undecidability is not available in the existing literature. Notably, verifying whether a given vector is an entropic vector is feasible only in specific cases, such as when the random variables involved are binary [35].

### 4.2 Linear approximation

In order to avoid the notably complex optimization problem (7), we shift from working directly with the space of probability distributions *q* to an entropic representation, where the variables *h*(*A*) represent the values of entropy H(**X**_*A*_), for each *A* ⊆ *N*. The collection of all such variables forms a polymatroid *h* and the entropic constraints in (7) further constrain the values *h*(*A*) of *h*. This leads to the following linear approximation of the initial problem (7).

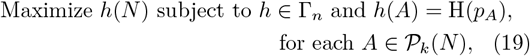

where the control vector variable is *h* ∈ ℝ ^*𝒫* (*N*)^. The feasible set of (19) consists of polymatroids *h* ∈ Γ_*n*_ such that the values of 2^*k*^ variables *h*(*A*) for |*A*| *≤ k* are fixed to H(*p*_*A*_). Additionally, for *n ≥* 4, the problem can be constrained further by incorporating all instances of the Zhang-Yeung inequality (18).

Conveniently, the linear program (19) is always feasible because the entropic vector *h*_*p*_, defined by (17), trivially satisfies the constraints. Moreover, the feasible set is bounded. Specifically, by the polymatroid constraints and the entropic constraints *h*(*i*) = H(*p*_*i*_) for *i* ∈ *N*, we can derive the following upper bound for any *A* ⊆ *N* with |*A*| *> k*:

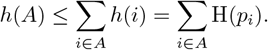

Although the formulation of linear approximation (19) doesn’t involve any condition on the sample space 𝒳, the constraints imply that

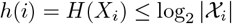

for each *i* ∈ *N*. This means that Γ_*n*_ includes only those entropic vectors (17) whose support size is less than or equal to the cardinality of the input sample space 𝒳_*i*_.

In conclusion, the optimal value of the linear program (19) serves as an upper bound for the optimal value of the original nonconvex problem (7), while the entropy of the input distribution H(*p*) provides a trivial lower bound. We summarize the properties of maximum entropy problems (4), (7), and (19) for a given input distribution of random vector (*X*_1_, …, *X*_*n*_) with the sample space × _*i*∈*N*_ *𝒳*_*I*_ and order *k ≤ n* in Table 5. Note that while (4) and (7) have the same number of variables, the former problem has at least as many constraints as the latter. The linear approximation (19) can have significantly less variables than either (4) and (7) since the dimension of problem (19) does not depend on the cardinality of the sample space. Most importantly, it concerns a simple linear optimization problem.

**Table 5:**
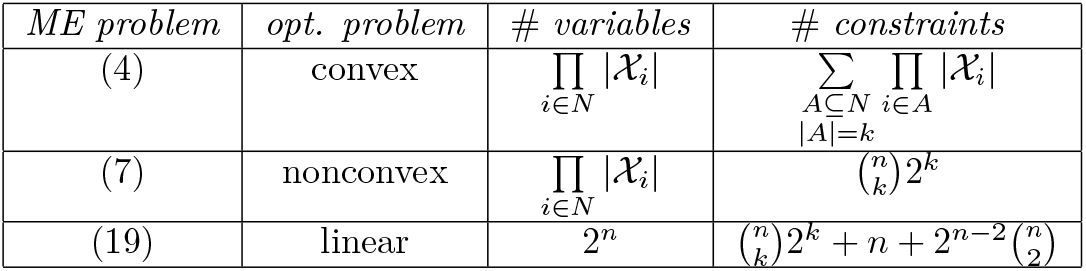
Comparison of ME problems.

### 5. Numerical experiments

Before illustrating the the proposed maximum entropy and connected information estimators on real-world datasets, we evaluate its properties in more realistic simulations than in the previous chapter. To this end we work with (discretization of) multivariate normal distributions. Normal distribution has the convenient property that its dependepce structure is fully described by its correlation (covariance) matrix, that is bivariate / second-order features, and thus in the maximum entropy/connected information sense it has no higher order interactions. More formally, we exploit the well-known fact that the *n*-dimensional normal distribution 𝒩 maximizes entropy among all distributions with a prescribed covariance matrix. Fixing the marginals of dimension 2 naturally also fixes the entire covariance matrix so that for a normal distribution 𝒩, and thus a given normal distribution is also the maximum entropy distribution with given second-order marginals. While we have strictly formally defined the connected information and the related optimization problem in the context of distributions on a finite set, its natural extension to distributions with continuous support thus provides 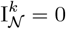 for all *k >* 2.

By continuity arguments one can thus expect that for suitably discretized approximation of normal distribution, the connected information estimates for *k >* 2 should also converge to 0 with decreasing discretization granularity (and increasing sample size in case of numerical sampling). On the other side, the entropic constraints are not as strict, as discussed in previous chapters, and thus we can expect the entropic connected information to be above zero even for *k >* 2. Conveniently, we can thus use the behaviour of connected information for mulrivariate normal distribution to evaluate the tightness of the optimization with entropic instead of marginal constraints.

The marginal constraint optimization requires the distribution to be defined on a finite set. To allow for a comparison between the two choices of constraint, we discretise the distribution via binning and estimate the CI from the sample distribution. While the binning distorts the normal distribution, we expect to see 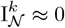 for *k >* 2 especially with larger numbers of bins. In Fig. 1, we can see indeed that when using marginal constraints, the values 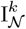 for all *k >* 2 are very close to zero. On the other hand, we show the presence of non-zero terms 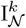 at all orders when using entropic constraints, although they decrease relatively fast above *k >* 2. This is in line with the expectation that entropic connected information provides a bound on connected information which is however not necessarily very tight.

**Figure 1.**
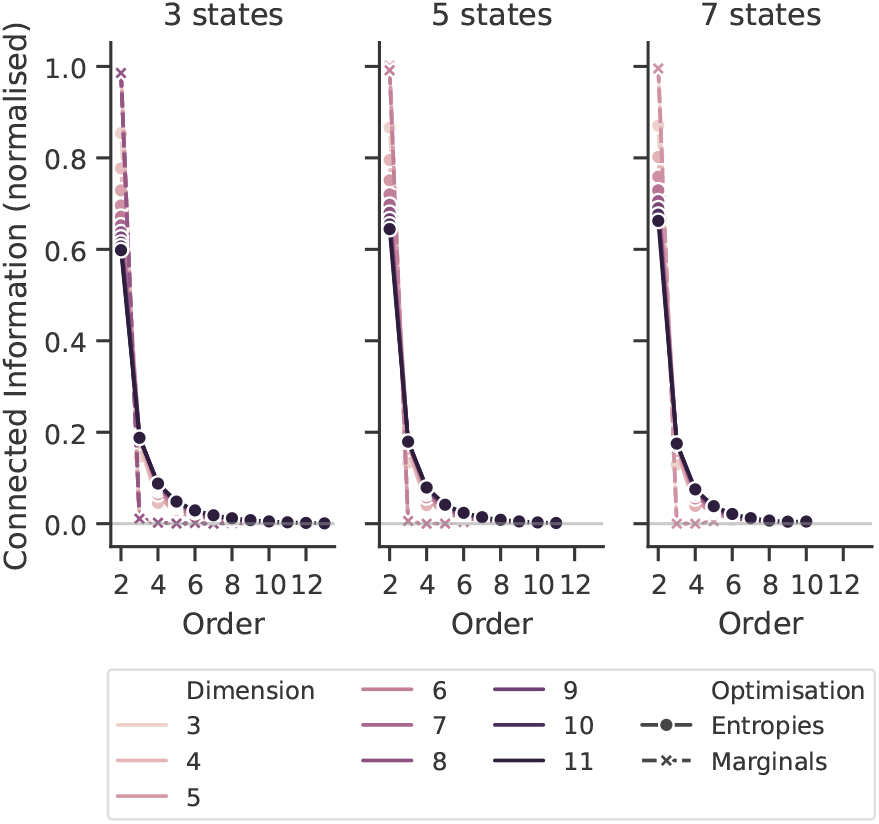
CI for multivariate normal distributions discretised into 3, 5, and 7 states. We show the results of 10 independent samplings (10^7^ samples) from the same multivariate normal distribution for each dimension.

Notably, the increased precision of the marginal constraint optimization comes with a considerable computational cost, preventing us from including the higher dimensionalities for the marginal constraints in Fig. 1. We show in Fig. 2A the running times of the two approaches on discretised multivariate normal distributions. While the complexity of both problems grows exponentially with the dimensionality, constraining the marginal distributions makes the computation much faster. We show in Fig. 2B how the exponential factor grows with the cardinality of the sample spaces | 𝒳_*i*_| (here assumed identical for all *i*). Notably, on the other hand, the factor is almost constant using the entropic constraint, showcasing the explosive computational advantage of the entropy-based connected information formulation for variables with high cardinality. One may wonder if the deviation from zero connected information is due to insufficient sampling or entropy estimation is affecting the optimization with the entropic constraints. To explore whether this is (not) the case, we take the limit of infinite samples and an infinite number of bins. It is possible to perform a similar analysis running the ‘example_ContinuousGaussian.jl’ script in the ‘examples’ folder in [32]. When fixing the marginal entropies, we can replace the sample estimate of the entropies with the analytic results for multivariate normal distributions. Fixing the marginal entropies discards the information on the covariance structure and eludes the constraint that was making the distribution maximally entropic. Thus the optimization thus gives an upper bound to the actual entropies 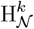. In Fig.3, we show the continued presence of non-zero terms 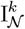 at all orders under entropic constraints, close to the numerical estimates provided for finite discretization and sampling. This demonstrates that the sampling based estimate of entropic connected information is reasonably precise, and its non-zeroness is not an sampling estimate artifact.

**Figure 2.**
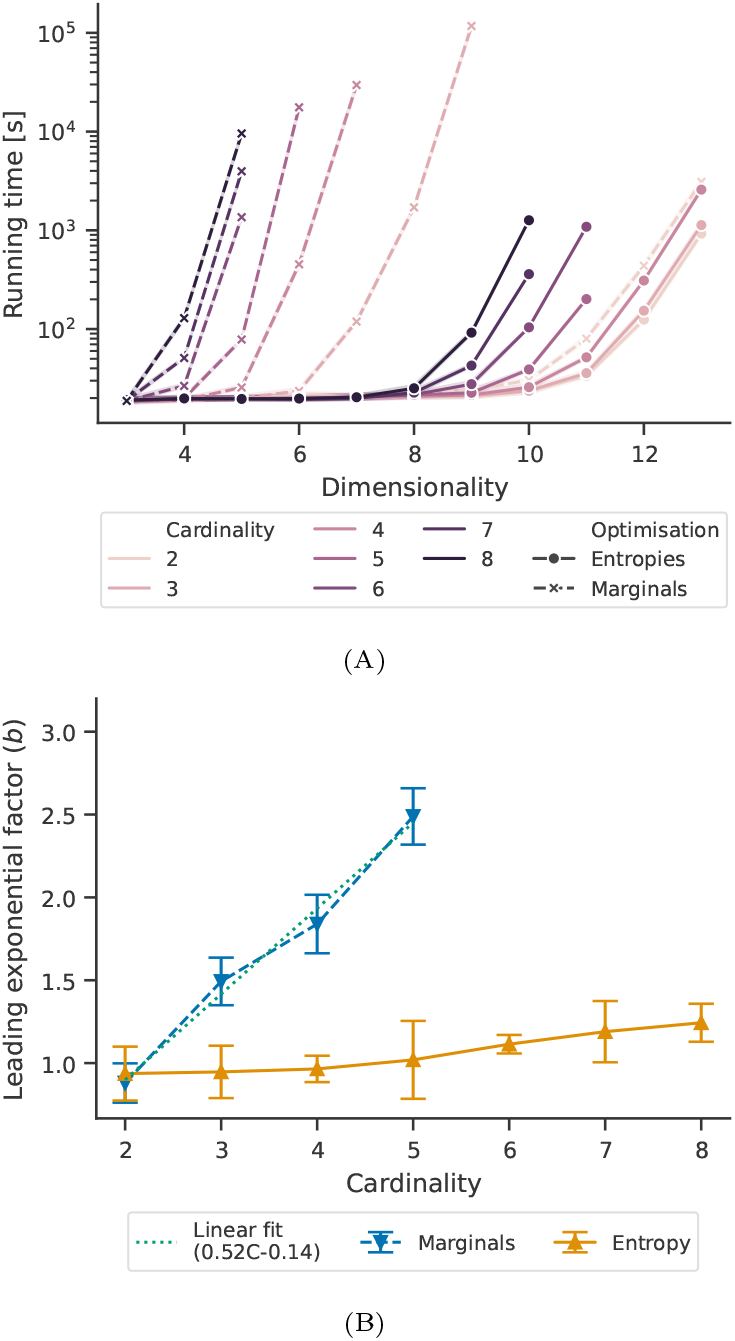
A) Benchmark of the optimization’s running time (in seconds) with constraints on the marginals or entropies. The running time was capped at 10^5^ ∼s. While the running times seem to grow faster than exponential, the best fit is given by a function like *t* = *const*. + *a*e^*bD*^ + *cD*^2^. B) Values of the coefficient *b*. In the optimization with fixed marginals, we show the fit parameters up to cardinality 5. With higher cardinality, the computational time is prohibitively long to obtain a sufficient number of data points for the fit.

**Figure 3.**
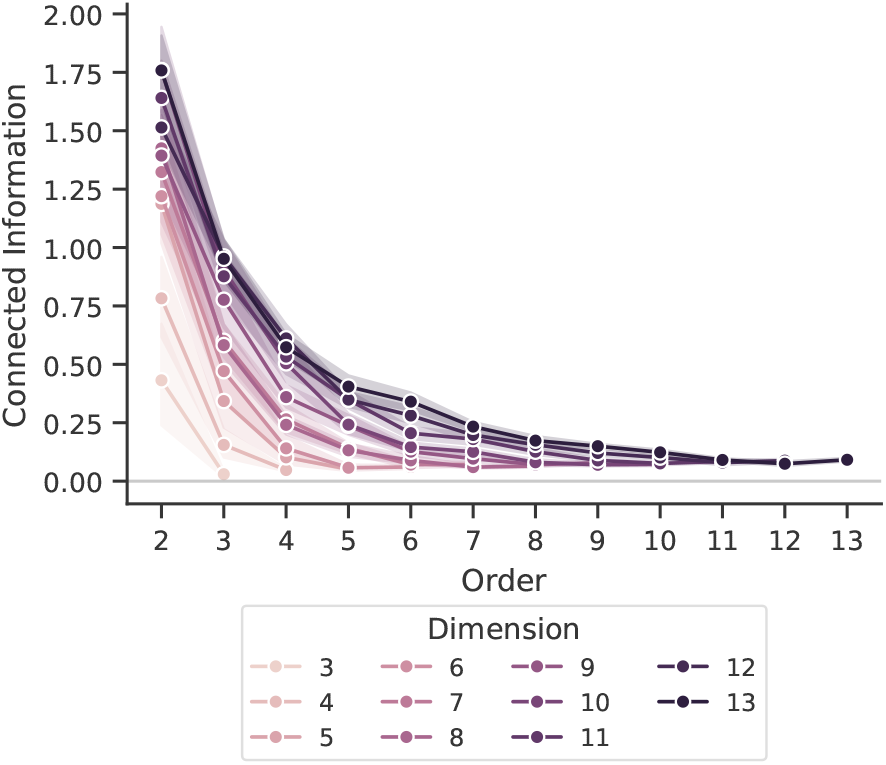
CI estimated with entropic constraints for multivariate normal distributions with 15 different random covariance matrices for each number of variables. Mean and range is shown by the line/shaded area.

### 6. Real-world examples

The entropic constraints enable the computation to be fast enough to study larger systems. However, the upper bound approximation might in principle be so rough that it would conceal any insight about the higher-order interactions among variables. We tested the maximisation with entropic constraint in realistic conditions to show that, beyond the approximation, key properties of the distributions are preserved. We chose three different systems: characters in texts, neuronal spikes, and DNA bases.

#### 6.1. Texts

The first system is written language, where we look at the connected information in the probability distribution of strings of subsequent characters in texts. It is possible to perform a similar analysis on custom data running the ‘example_Texts.jl’ script in the ‘examples’ folder in [32]. The random vectors **X** are character *n*-grams, sequences of *n* consecutive characters occurring in a text. We obtain our empirical distribution as the frequency of the *n*-grams over all the *n*-grams in a large corpus.

The *X*_*i*_, *i* 1 ∈, …, *n* take values over all possible Unicode points. To reduce the space 𝒳 and ensure that our estimate of the probability is not affected by undersampling for larger *n*, we grouped the characters in three classes according to their frequency: high, medium, and low, such that an equal number of characters in the text belongs to each class.

The nature of the *n*-grams as non-independent substrings of the original text allows us to check if the connected information at higher orders is reasonable. We fitted a *k*th-order Markov Model (*k < n*) to our corpus, and we used the model to produce a new stream of character classes from which we estimate a new probability distribution. The connected information should follow the one observed in the actual text up to order *k* + 1 and then drop to the noise level.

Figure 4 reports the results for *n* = 6 and *k* ∈ {1, 3, 5}. We repeated the analysis in two languages: English, considering all the novels and the 100 most prolific blog authors from [34] for a total of 2.35 *×*10^8^ *n*-grams, and German, using one million sentences in the 2023 news dataset of the Leipzig Corpora Collection [18] containing 1.04 *×* 10^8^ *n*-grams.

**Figure 4.**
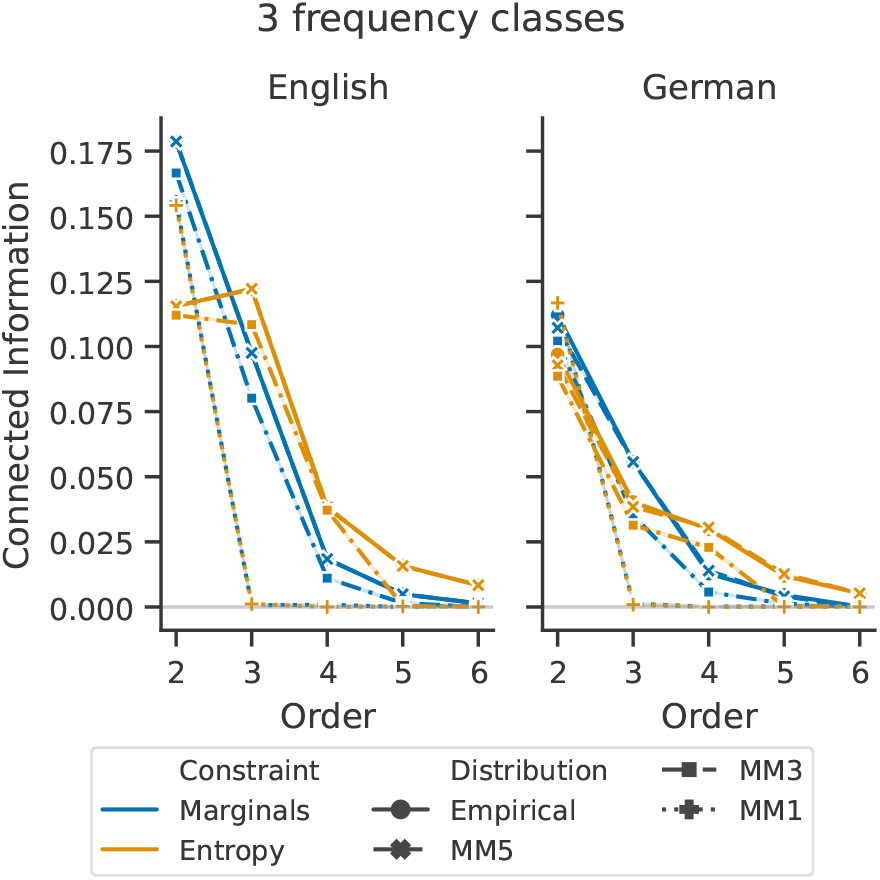
Connected information in the character probability distribution in English (left) and German (right) example text corpora. Different colours represent different constraints in the maximisation of the entropy. Solid lines represent the empirical distribution, while dashed lines represent its approximation through Markov Models of a different order (MM1: Markov Model of order 1, and so on). The drop in connected information for the Markov Models of order *k* correctly reflect their inability to capture relationships beyond the order *k* + 1, in particular, the drop in connected information for the Markov Model of order 3 shows its inability to capture relationships beyond order 4, while the Markov Model of order 5 reproduces the dependency structure for all six orders. Both the Marginal-based and Entropy-based CI estimate suggest that empirical data contain higher order interactions that the Markov Model truncated. Note that the optimization with fixed entropies yields slightly higher values of CI than optimization with fixed marginals for higher orders, which is expected due to the upper-bound nature of the entropy-based approximation.

If, instead of three character classes, we assign each letter to its state (and add a 27th state for non-alphabetical characters), we are already far beyond the treatable regime for the fixed marginal optimisation. Conversely, the optimisation problem is solved in minutes with the entropic constraints. While we know this is just an approximation of the CI for the distribution, we can still appreciate the difference in Fig. 5 with the Markov Chain surrogates. The difference between the empirical distribution and the MM order 2 for 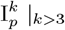 clearly shows the relevance of higherorder interactions in this system.

**Figure 5.**
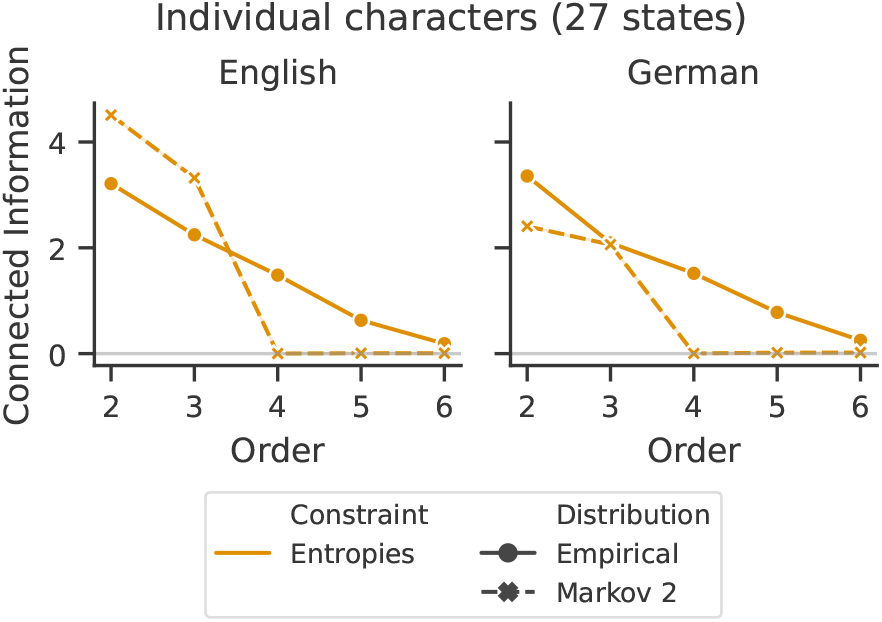
Connected information (entropy-based approximation) in the character probability distribution in English (left) and German (right) example text corpora. Different colours represent different estimates of the entropy. Solid lines represent the empirical distribution, while dashed lines represent its approximation through a Markov Model of order 2. The drop in connected information for the Markov Model shows its inability to capture relationships beyond order 3. This property is conserved for many states (and an undersampled distribution) despite the computation of upper bounds.

We do not expect the CI in the Markov model at orders 2 and 3 to match exactly the estimate on empirical data. Indeed, the Markov model reproduces only some of the order 2 and 3 projections of the 6-dimensional distribution. Moreover, we point out that this distribution is heavily undersampled, with ∼1.6 states for every sample. As we show in the following example, this can severely impact the estimates and we introduce a possible way to alleviate the problem.

#### 6.2. Neuronal dynamics

The second system we consider is the neuronal activity in the mouse brain. In this case, we consider individual units and their instantaneous spiking rate. We used the spiking times from all the mice (N=26) that underwent the Functional Connectivity experiment from the Allen Brain Observatory – Neuropixels Visual Coding dataset [30] which includes 1800 s of recording in resting condition.

In this scenario the undersampling of the high-dimensional distribution is one of the limiting factors. Estimating the firing rate over fixed intervals, long enough – at least 100 ms – to ensure adequate sampling even for sparsely firing units, would yield a relatively small number of samples. Instead, we estimate the instantaneous firing rate for each unit every time any unit fires. We consider a window equal to the unit’s average interspike interval and check whether it contains zero, one, or more spikes, corresponding to three different classes. This procedure provides ∼100 samples per second, sufficient for sampling up to dimension seven. We thus selected two groups of 7 units from each session based on their quality metrics.

Figure 6 reports the connected information in the system. To compare the estimated interactions in the empirical data to a system with known, lower-order structure, we generate specific surrogates with destroyed spatial structure by shifting together the spiking times of selected groups of units, destroying the interactions of orders above the group size, while preserving at least part of the lower orders. Notice that the basic plug-in entropy estimate provides a spurious observation of a non-zero (and growing) estimate of CI even in the case of surrogates (where any higher-order interactions were purposefully destroyed by shuffling). We apply a correction due to Grassberger [19] to estimate marginal entropies and limit this effect. The correction prevents the unnatural growth of the CI estimate with increasing order (green curves in Fig. 6).

**Figure 6.**
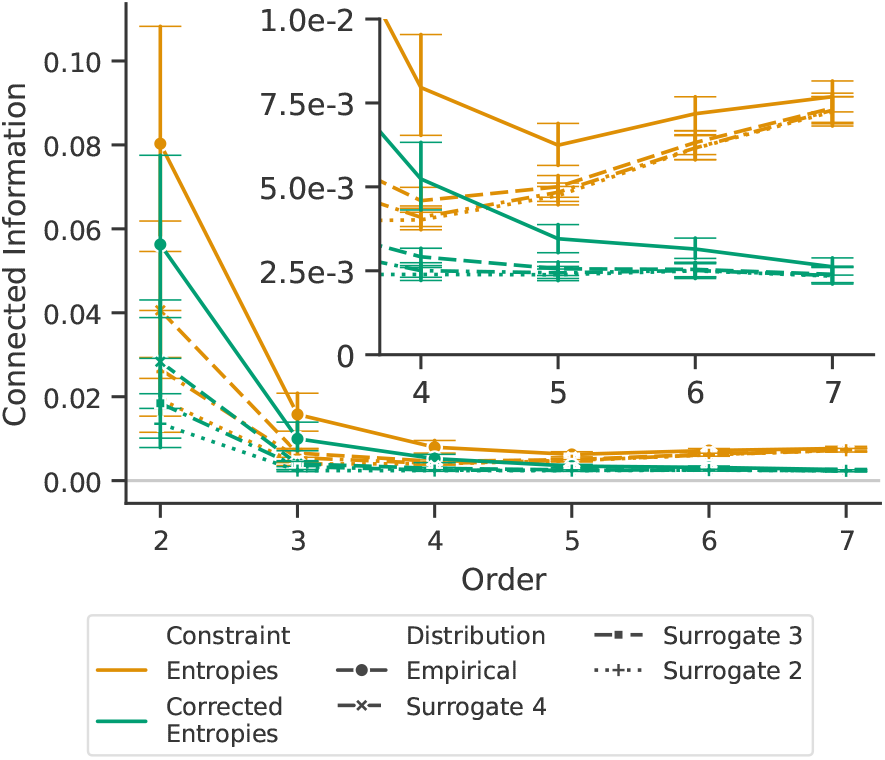
Average over mice and groups of units of the connected information in individual units firing rate for a meso-dimensional system (*N* = 7). The colour indicates the type of entropic constraint: raw or with the Grassberger correction. The number of possible states of the system is 2183 with a number of samples between 1.39 *×* 10^5^ and 2.23 *×* 10^5^.

Studying the CI behaviour in the simulations (surrogates), we observed that the CI estimate seems to plateau at about 2.5*×* 10^−3^ (which we can consider residual estimate bias). However, different surrogates approach this limit at different orders: when shifting pairs of units (i.e., Surrogate 2), the CI is flat from order three; when shifting four units, the curve reaches the plateau only at order five, with the plateau appearance corresponding exactly to the order where interactions were precluded by the shuffling. We can thus establish that HO interactions in the empirical data remain above those of surrogates at least up to the seventh order.

We can wonder whether this convergence towards the surrogates of the CI estimated from the empirical distribution is visible beyond the system dimension for which the Grassberger’s correction provides reasonably unbiased information estimates. Indeed, it could be an artefact due to the chosen dimensionality. Thus, we consider the distribution of the 12 highest quality units for each mouse. This is now clearly a prohibitively large dimension to solve with marginal constraints. Although the optimization itself with entropy constraints is still viable in terms of computational demands, even Grassberger’s correction for sparse sampling is now insufficient to compensate for the bias in CI estimates due to undersampling effects above order 6. Nevertheless, in terms of statistical inference, we observe in Fig. 7 that the CI estimated from the empirical distribution stays above the surrogates until order 7 (and becomes similar or lower onwards). This information can inform subsequent analysis of the system by limiting the highest-order terms worth further study with in-depth tools.

**Figure 7.**
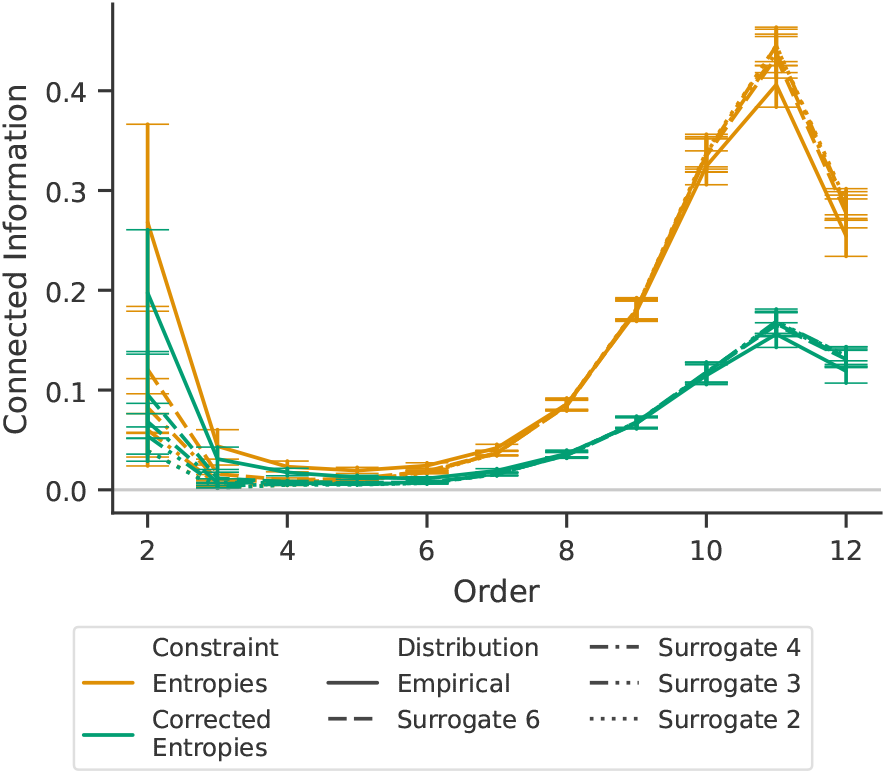
Average over mice and groups of units of the connected information in individual units firing rate for a high-dimensional system (*N* = 12). The colour indicates the type of entropic constraint: raw or with the Grassberger correction. The distribution is undersampled, with 5.32 *×* 10^5^ possible system states and a sample count between 3.05 *×* 10^5^ and 4.74 *×* 10^5^. In the inset, we show the difference between the CI estimate based on the empirical distribution and that based on the surrogates.

So far, we have shown that optimisation with entropic constraints preserves a significant difference between the empirical distributions and distributions with known zero CI at specific orders. In the next example, we show how the optimisation can confirm expected differences between empirical distributions.

#### 6.3. Genomics

As a last example, we test the connected information with entropic constraints in another biomedical context. It is possible to perform a similar analysis following the instructions in the ‘example_DNA.jl’ script in the ‘examples’ folder in [32].

Eukaryotic genomes consist of coding and non-coding DNA sequences. A notable difference is codons, 64 specific 3-nucleotide-long sequences that encode individual amino acids in the coding DNA. The codons include all possible three-base sequences, and their sequences cannot be immediately distinguished from those of non-coding DNA. However, their relationships, as reflected in coding DNA, mirror those between amino acids in proteins. The presence of codons in coding DNA introduces interactions at short to intermediate orders. In particular, we expect to observe a significant difference in the CI of coding and non-coding sequences at orders 4-6, corresponding to pairs of codons. We acknowledge that this is a crude simplification of DNA’s structure. For example, we gather in the same group non-coding sequences that include introns, untranslated regions at the extremities of genes, regulatory elements, and noncoding genes. In the latter ones, some dependencies are likely introduced by RNA secondary structures, analogous to those introduced by protein secondary structures in coding DNA. These can span tens of bases and introduce long-range interactions. However, in both coding and noncoding DNA, we expect the CI to gradually decrease at higher orders due to heterogeneity in these higher-order interactions across the genome.

To test whether we can detect interactions between codons in DNA sequences, we gathered DNA sequences from 250 organisms. These are all the species available in the Ensembl database [14] available at [15] with at least 10M bases of protein-coding DNA, excluding duplicates such as *C. lupus familiaris, C. lupus familiaris basenji, C. lupus familiaris boxer*. This dataset contains a wide variety of animals including mammals (*H. Sapiens, Bos taurus, Felis catus*), birds (*Falco tinnunculus, Anas platyrhynchos, Phasianus colchicus*), fish (*Electrophorus electricus, Sparus aurata, Salmo salar*), reptiles (*Pogona vitticeps, Pelusios castaneus, Anolis carolinensis*), and invertebrates (*Drosophila melanogaster, Caenorhabditis elegans*).

We downloaded the full genome (*.dna.toplevel.fa) and subtracted the coding sequences (*.cds.all.fa) to obtain the noncoding DNA. We computed the probabilities of 8-base sequences separately for each organism and DNA type.

In Figure 8 we show the connected information at orders 2 to 8. The significantly higher CI in coding sequences at orders 4 and 5 aligns with our hypothesis based on the patterns in coding DNA. The higher CI for noncoding sequences at orders 7 and 8 warrants further investigation.

**Figure 8.**
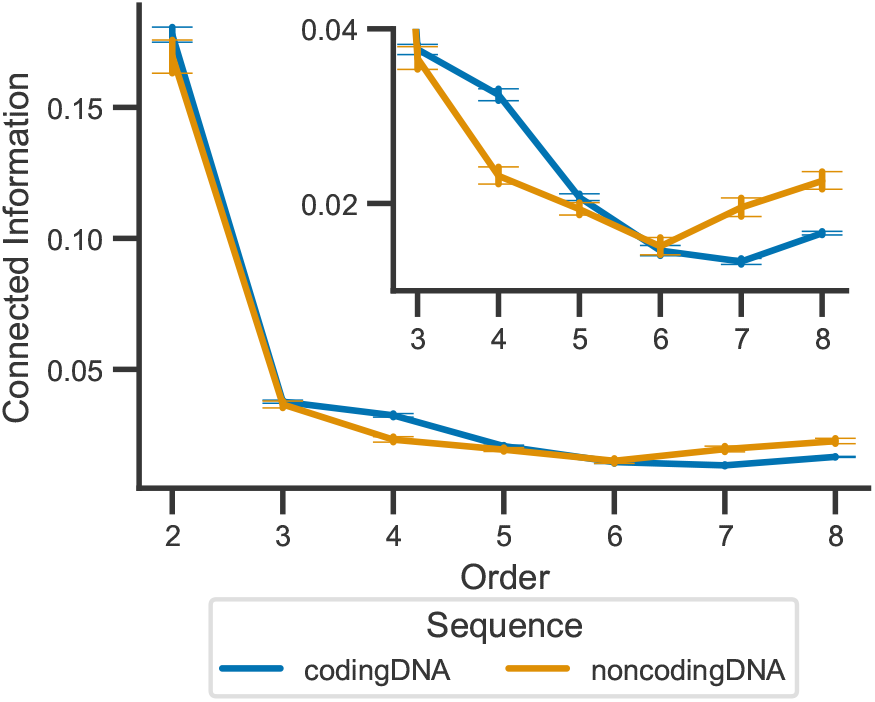
Average over 250 species of the Connected Information in coding and noncoding DNA. The errorbars mark the 95%c.i. At orders 4, 5, 7, and 8, the CI in the two kinds of DNA is significantly different (*p <* 0.01, Bonferroni corrected).

A possible source is an effective undersampling of distributions due to the high repetitiveness of some non-coding sequences, which would drive an apparent growth of the CI for high orders as observed in Fig. 7. However, as the organisms that drive the difference have longer non-coding DNAs, this may reflect true information, e.g., as introduced by the secondary structure in long non-coding RNA [21]. Testing this hypothesis is not possible with the current dataset: selecting only non-coding RNA sequences to corroborate the secondary structure as a source of true CI forces the use of very short sequences, introducing biases due to undersampling.

## 7. Discussion and conclusion

We presented a general framework for estimating connected information from maximum entropy problems constrained by marginal or entropic quantities, together with its practical implementation in the HORDCOIN library. This formulation extends classical moment- and marginal- based approaches by providing a theoretically grounded and computationally tractable route to quantifying higher-order statistical dependencies in multivariate data.

The proposed linear programming approximation offers a practical balance between analytical rigor and scalability. Our results show that, despite its upper-bound nature, the entropy-constrained formulation captures the essential structure of higher-order interactions in both synthetic and empirical systems. In particular, the experiments on language data confirmed the expected decay of connected information beyond the order of dependence captured by Markov models, while analyses of neuronal population activity revealed significant higher-order dependencies persisting up to the seventh order. The findings demonstrate that the entropic formulation remains robust even under strong undersampling that typically hampers classical estimators. Beyond methodological innovation, the proposed framework opens the way for systematic characterization of higher-order dependencies in large-scale neurophysiological and neuroimaging data. By providing a fast, open-source implementation, HORDCOIN lowers the technical barriers to applying information-theoretic tools in biomedical contexts. Future developments may integrate bias-correction techniques and hybrid entropic–marginal constraints to further improve accuracy in small-sample regimes and enable the study of continuous-valued variables.

In summary, the entropic-constraint approach to connected information provides a theoretically elegant, computationally efficient, and practically useful tool for disentangling multivariate dependencies in complex biological systems, bridging the gap between abstract information theory and real-world biomedical data analysis.

## CRediT authorship contribution statement

**Giulio Tani Raffaelli:** Data curation, Formal analysis, Investigation, Methodology, Resources, Software, Validation, Visualization, Writing – original draft, Writing – review & editing.

**Jakub Kislinger:** Formal analysis, Investigation, Methodology, Software, Validation, Writing – review & editing.

**Tomáš Kroupa:** Conceptualization, Formal analysis, Funding acquisition, Investigation, Methodology, Project administration, Resources, Supervision, Writing – original draft, Writing – review & editing.

**Jaroslav Hlinka:** Conceptualization, Formal analysis, Funding acquisition, Investigation, Methodology, Project administration, Resources, Supervision, Writing – original draft, Writing – review & editing.

**Acknowledgements**

The authors were supported by the Czech Science Foundation project No. 21-17211S and by the Johannes Amos Comenius Programme (P JAC) provided by MSMT (reg. n. CZ.02.01.01/00/23_025/0008715. The authors thank Marika Kosohorská (Czech Technical University in Prague) for helpful comments on an earlier draft of this paper. The authors thank Julia Steger for the precious advice on the data selection and interpretation for the DNA example.

## 8. Declarations

### 8.1. Conflicts of interest

The authors have no conflicts of interest to declare that are relevant to the content of this article.

### 8.2. Data availability statement

This study used publicly available datasets: language [34, 18], neuronal data [30], genetic data [14]. The code is available at [32] and [33].

## Notes

https://github.com/cobragroup/Hordcoin.jl

https://github.com/cobragroup/pyHordcoin

### Competing Interest Statement

The authors have declared no competing interest.

